# No general shift in spring migration phenology by eastern North American birds since 1970

**DOI:** 10.1101/2021.05.25.445655

**Authors:** André Desrochers, Andra Florea, Pierre-Alexandre Dumas

**Affiliations:** Département des sciences du bois et de la forêt, Université Laval, Québec, Canada; Observatoire d’oiseaux de Tadoussac, Québec, Canada; Observatoire d’oiseaux de Tadoussac, Québec, Canada and Département des sciences du bois et de la forêt, Université Laval, Québec, Canada

**Keywords:** Birds, eBird, Migration, Climate change, Phenology, North America

## Abstract

We studied the phenology of spring bird migration from eBird and ÉPOQ checklist programs South of 49°N in the province of Quebec, Canada, between 1970 and 2020. 152 species were grouped into Arctic, long-distance, and short-distance migrants. Among those species, 75 significantly changed their migration dates, after accounting for temporal variability in observation effort, species abundance, and latitude. But in contrast to most studies on the subject, we found no general advance in spring migration dates, with 36 species advancing and 39 species delaying their migration. Several early-migrant species associated to open water advanced their spring migration, possibly due to decreasing early-spring ice cover in the Great Lakes and the St-Lawrence river since 1970. Arctic breeders and short-distance migrants advanced their first arrival dates more than long-distance migrants not breeding in the arctic. However, there was no difference among migrant groups when median arrival dates were considered. We conclude that general claims about advances in spring migration dates in eastern North America are misleading due to large taxonomic variation.

## Introduction

Spring migration in birds is triggered by a combination of physiological, astronomical, and climate factors (Gwinner, 2003; Newton, 2008). Astronomical and physiological phenomena do not vary at the decadal scale, but climate and weather do, with substantial variation at the regional level (IPCC, 2013). With the rising awareness of climate change, ornithologists have focused on its possible effects on avian ecology, especially migration phenology (Berthold, 1991), ensuing ecological mismatches (Both and Visser, 2001; Thomas et al., 2001; Saino et al., 2009), and their possible consequences on populations (Møller et al., 2008; Both et al., 2006). Warmer weather is generally thought to advance spring migration, e.g. by reducing the risk of late frosts and increasing food availability early in the season due to changing plant and arthropod phenology (Gordo, 2007). As a result of the accumulating studies on the subject, the Intergovernmental Panel on Climate Change (IPCC) gave “high confidence” in the claim about a general spring advancement of migration in the northern hemisphere (Settele et al., 2014). Some authors went further and labeled this phenomenon as a ‘flagship example of the biological impacts of climate change’ (Kelly et al., 2017), and one of the best documented biological responses to climate change (e.g. Miller-Rushing et al., 2008; Newson et al., 2016).

Early evidence for the advancement of spring migration came mostly from numerous small-scale studies, conducted mostly in Europe (Gienapp et al., 2007; Leech and Crick, 2007). A close examination of those studies shows that the changes in spring migration dates are not as straight-forward as implied by the above claims. For example, Lehikoinen (2004) found that of 222 time series, only 26 % exhibited significantly advancing mean migration dates. Most time series (69 %) exhibited no pattern and 5 % showed delayed migration. A more recent review based on 440 time series of mean or median arrival dates for 214 species also suggested a great variability in trends, with evidence for an advancement (28 % of the species), no change (51 %) and a delay (21 %) in spring migration dates [Lehikoinen and Sparks (2010); Fig. 9.4]. Knudsen et al. (2011) reviewed ten inferences made about phenological patterns in migration from recent decades, and Chmura et al. (2019) provided a detailed review of mechanistic hypotheses for those patterns. Most of these studies associated phenological changes to the warming climate of recent decades, because the former “is consistent with” the latter. But empirical evidence for this causal inference is often questionable, due to the presence of several sources of variation such as species life-history, geographic region, and age distribution (Knudsen et al., 2011). Knudsen et al. (2011) point out that because species differ in their movements and ecology, and because climate change varies geographically, we should not expect changes in migration phenology to be globally similar, but to be associated to regions as well as aspects of species ecology such as the location of their wintering grounds (Hüppop and Hüppop, 2011).

Despite the large number of migration phenology studies, empirical gaps in our understanding of regional variation still greatly limit our ability to understand spatio-temporal patterns, not to mention their causes and consequences. Those gaps do not come only from uneven sampling effort, but also from methodological misconceptions. Temporal changes in sampling effort (Miller-Rushing et al., 2008; Moussus et al., 2010) or bird abundance (Miller-Rushing et al., 2008; Tryjanowski et al., 2005; Sparks, 1999; Koleek et al., 2020) are known to bias trend estimates. This problem has been recognized by several authors (e.g., Moussus et al., 2010; Gordo, 2007; Goodenough et al., 2015), especially when extreme values such as first arrivals or low quantiles are used. The latter studies recognize the advantages of mean or median arrival dates, less sensitive to sampling effort and population size. Nevertheless, first, or early arrival dates continue to dominate the empirical basis for multi-decadal changes in spring migration (Tryjanowski et al., 2005; Knudsen et al., 2011). Despite its inherent sensitivity to sampling bias, abundance and the cumulative distribution of arrivals (Sparks et al., 2001), first arrival dates are worth examining in addition to central estimates such as medians and means, because changing strategies of early individuals may inform us about the plasticity of the species to changing environmental conditions (Mathot et al., 2012).

Another problem is the lack of reliable multi-decadal records outside those obtained from standardized migration counts at long-established bird observatories, mostly located in Europe. The advent of citizen science programs such as eBird (Sullivan et al., 2009) enable us to monitor changes in bird behaviour and populations in seasons usually not covered by standardised long-term programs such as the Breeding Bird Survey (Sauer et al., 2014) and the Christmas Bird Count (Butcher et al., 1990). Furthermore, citizen science offers a wider geographic coverage than previously available. As a result, it is possible to examine relationships between spring arrival dates in the North, and departure dates from wintering areas.

Here, we investigate whether timing of avian spring migration changed over the last 51 years (1970-2020), in Quebec, Canada, based on eBird (Sullivan et al., 2009) and *Étude des populations d’oiseaux du Québec* (ÉPOQ; Cyr and Larivée, 1979, 1993) checklist programs. We also test the relationship between migration phenology and migration distances, the latter obtained from January and February eBird records of each species. Finally, we assess whether variation in arrival dates of early species is greater than that of later-arriving species.

## Methods

We downloaded data from the February 2021 release of the eBird (eBird Basic Dataset, 2021) and ÉPOQ (Cyr and Larivée, 1979, 1993) datasets. Together, those datasets totalled 3,933,296 records for retained species between springs 1970 and 2020, between 1 March and 10 June. Since the launch of eBird, annual reported effort by Quebec birders has increased exponentially from 2,571 h (1970) to 114,840 h (2020; Fig. 1).

**Figure 1:**
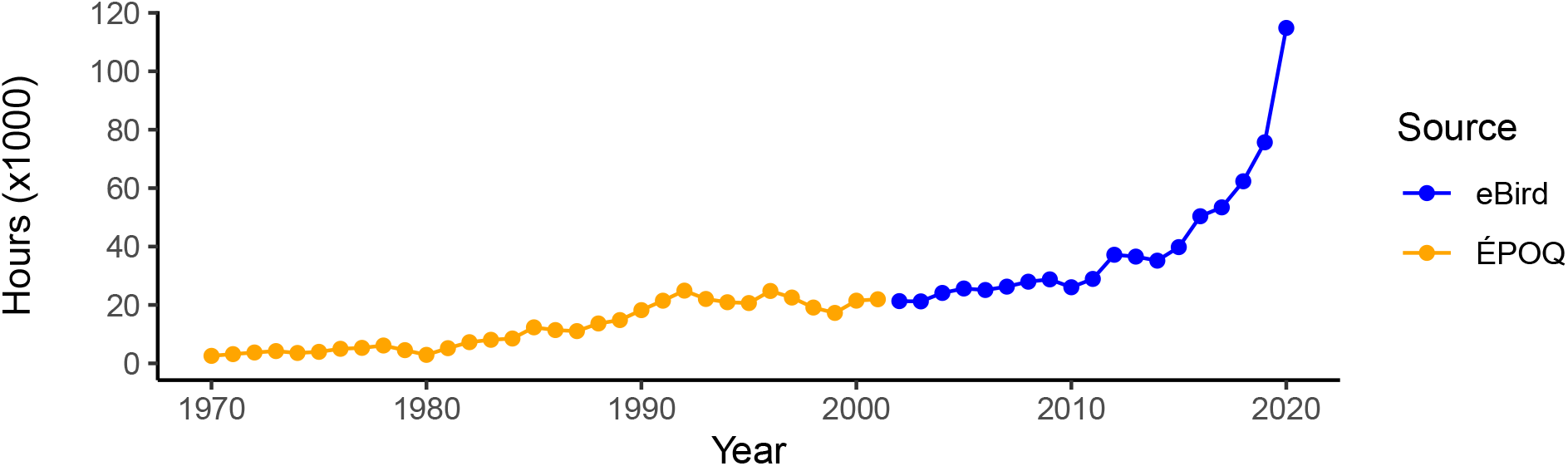
Observation effort (1 March - 10 June) from ÉPOQ and eBird checklist programs, south of 49°N, Quebec, Canada.

### Data preparation

We omitted data before 1 January 1970 from the analysis, because of small sample sizes. We also omitted data North of 49°N to reduce statistical noise due to latitudinal effects on bird arrival dates. Finally, we excluded all records before 1 March or after 10 June to reduce the influence of records of wintering or breeding birds. We selected species that were found in > 0.1 % of complete eBird lists, and whose records between 15 December and 15 February represented < 1 % of the species’ records.

We classified species into three groups based on migration distance: short-distance, long-distance, and Arctic migrants (Appendix). For short- and long- distance migrants, we extracted all midwinter (January and February) eBird records, between longitudes −100 and −30, i.e. covering Eastern North America and Latin America. We calculated maximum daily counts for each unique combination of species, latitude and longitude (1-degree squares), and date, and calculated mean midwinter coordinates weighted by the sum of daily counts. Short-distance migrants were species whose majority of individuals recorded in midwinter were north of 28°N, roughly corresponding to the north of the Gulf of Mexico. The other species were classified as long-distance migrants, except species nesting mostly in the Arctic, regardless of their wintering grounds. We made this distinction because the Arctic biomes are regarded as the most affected by climate change (IPCC, 2013). Thus, Arctic breeding species could be more affected by climate change than short or long-distance migrants.

Several indicators can be used to measure bird migration phenology, each with its own caveats (Moussus et al., 2010). We used two indicators: the first arrival date, and the median arrival date. For first arrival dates, each year we randomly sampled, without replacement, a number of records equal to the lowest number of records in the time series (min. 10) to account for changes in sampling effort and bird abundances, both affecting the number of records and in turn, the records’ earliest dates. Each year, we retained the earliest date from the random sample, as per Francoeur et al. (2012). We repeated the latter sampling procedure 100 times for each species, and computed the mean of the 100 first arrival date resamples for each year. For median arrival dates, we used Spark’s ‘median bird’ method (Bulmer, 1980), i.e. in the present case, the date at which half of the individuals recorded for the 1 March - 10 June date range were observed. Multiple checklists at the same time and location could duplicate the number of birds. Thus, we aggregated records by locality, date, and species, and retained maximum counts. Besides annual changes in sample size and species abundance, first arrival and median arrival dates could still be biased by annual variation in the timing of observation effort (e.g., early vs. late spring). Thus, we calculated the mean date of checklists each year to measure and account for this potential bias.

### Statistical modeling

We performed the analysis in two steps. First, we generated regression estimates for each species, and second we performed a meta-analysis of the regression estimates across species. For the first step, we used simple Gaussian linear models of arrival dates for each species as a function of year, annual mean effort (checklist) date, and mean checklist latitude as fixed effects. Effort date and latitude were included because of possible biases due to systematic changes in observation dates and locations that may have occurred in the 51 years of the study.

For the second step we tested whether mean regression estimates from step one differed from zero and whether they were associated with migration distance. There may be a substantial phylogenetic effects in migration phenologies (Rubolini et al., 2007). Thus, we used phylogenetically independent linear contrasts (Freckleton et al., 2002; Harvey and Pagel, 1991; Felsenstein, 1985) to account for phylogenetic proximity among the retained species, based on data from Jetz et al. (2012). Because of uncertain relationships between DNA data and years since speciation, phylogenetic trees are only approximations based on assumptions about the rate of phylogenetic divergence. Thus, we generated 100 phylogenetic trees from Jetz et al. (2014) and established phylogenetic distances for each dyad from the 152 species studied (Paradis and Schliep, 2018). We modelled the effects of migratory distance on regression estimates with a linear model using phylogenetic Generalized Least Squares (pGLS). The pGLS method accounts for the phylogenetic distance between species by using a distance covariance matrix which gives more weight to differences between phylogenetically distant species than differences between closely related species (Felsenstein, 1985; Grafen, 1989; Harvey and Pagel, 1991). We ran 100 pGLS models, corresponding to each of the 100 phylogenetic trees of all species. We calculated mean estimates, their standard error and their p-values from the 100 models. We used an error rate of *α* = 0.05 for significance statements, with no correction for multiple tests. All data preparation and statistical analyses were performed with R (R Core Team, 2020), with packages ape (Paradis and Schliep, 2018), `phylolm` (Ho and Ané, 2014) for pGLS, and `emmeans` for gls multiple comparisons (Lenth, 2021).

## Results

Since 1970, spring observers have been relatively constant in the dates of their observations, with a mean spring effort date advancing by −1.1 ± 0.7 days over the 51-y period (*p* = 0.1). After accounting for possible biases due to changing effort dates and latitudes, 75 species significantly changed their spring migration phenology, in terms of first arrival dates (48 spp.) or median arrival dates (52 spp.; see Appendix). However, first or median dates did not generally advance since 1970, (Fig. 2; 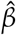 = −0.041 ± 0.24 days per decade, *t* = −0.16, df = 148, *p* > 0.9). When first and median dates are combined, 36 species advanced their migration by at least one metric, while 39 delayed their migration.

**Figure 2:**
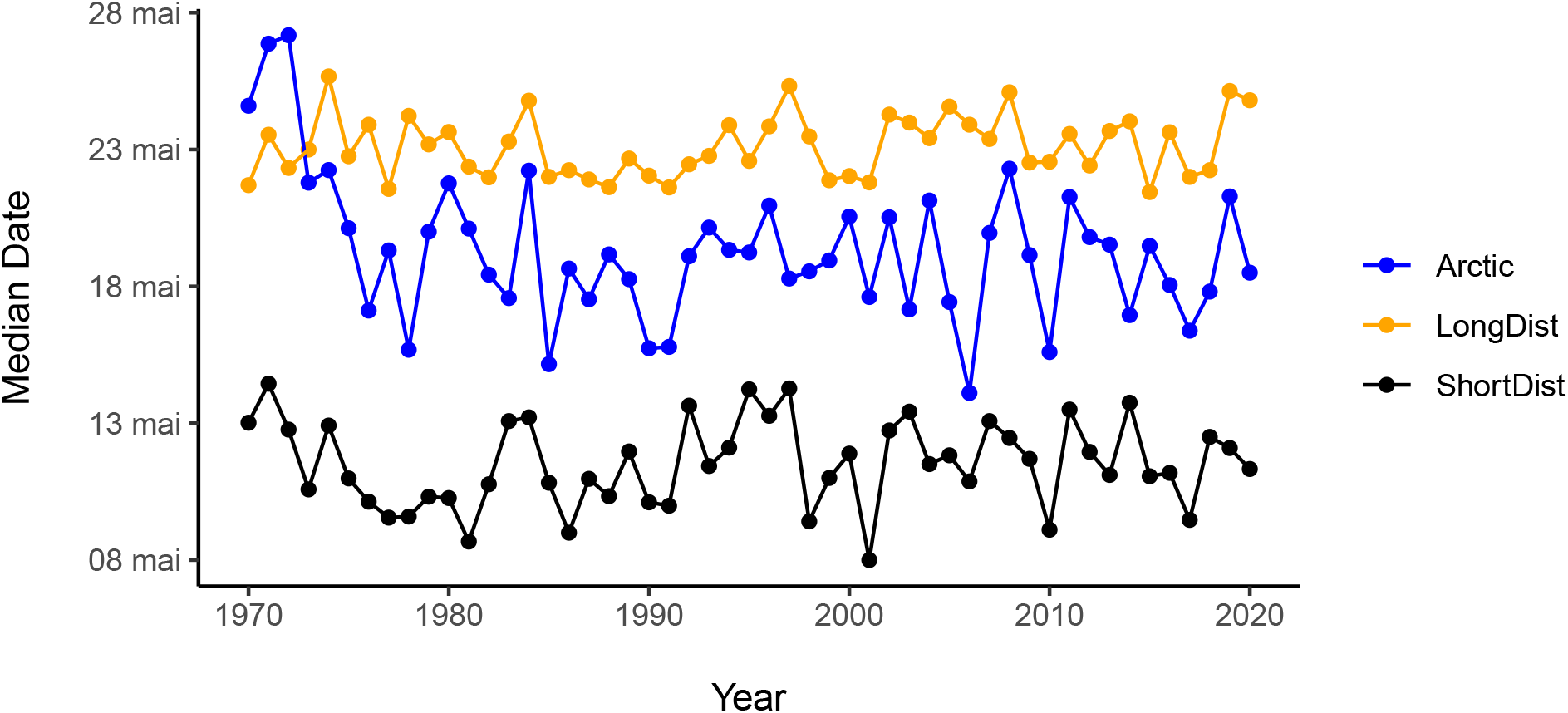
Changes in spring median dates according to migratory distance. Median dates were obtained by averaging species-specific dates at which 50 % of the individuals were recorded.

Based on median dates, 23 of the 149 species investigated showed a significant advance in spring migration, 29 species arrived significantly later since 1970, and 97 did not exhibit change. There was no significant variation of trends among species groups defined by migration distances (*F* = 0.08, df = 2, 146, *p* = 0.93). However, it is worth mentioning that the majority of the species that did advance their median arrival date were associated to open water whereas very few (~5) of the later-arriving species fell into this category (Fig. 3).

**Figure 3:**
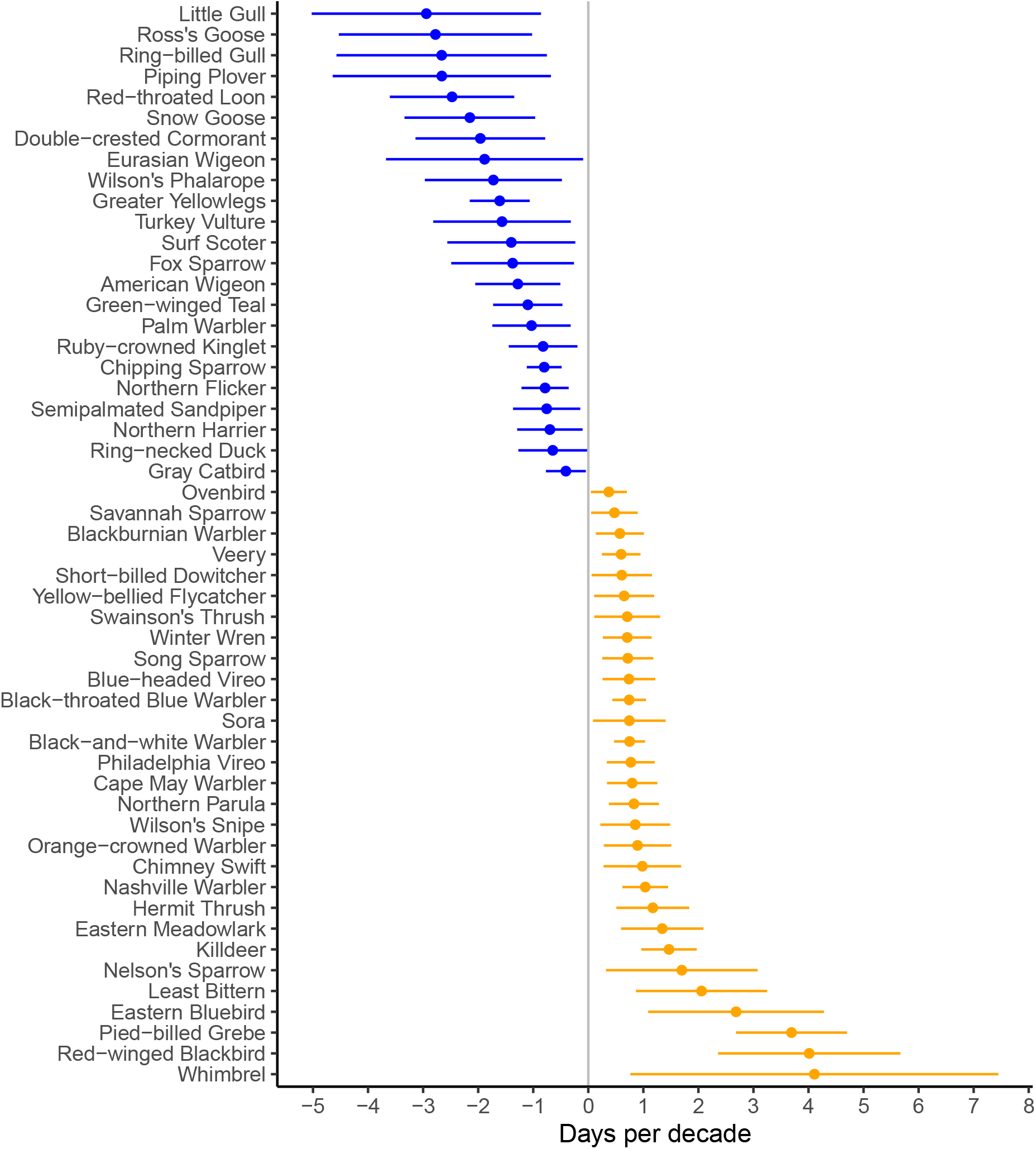
Species with significant 50-year trend in median arrival dates. Bars represent 95 % confidence intervals.

Similarly to median dates, first arrival dates did not generally advance since 1970, after accounting for bias in checklist dates and latitudes (Fig. 4; 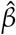 = −0.35 ± 0.46 days per decade, *t* = −0.77, df = 133, *p* = 0.4). 29 of the 134 species investigated showed a significant advance in spring migration, 19 arrived significantly later since 1970, and 86 did not exhibit change in first arrival date.

**Figure 4:**
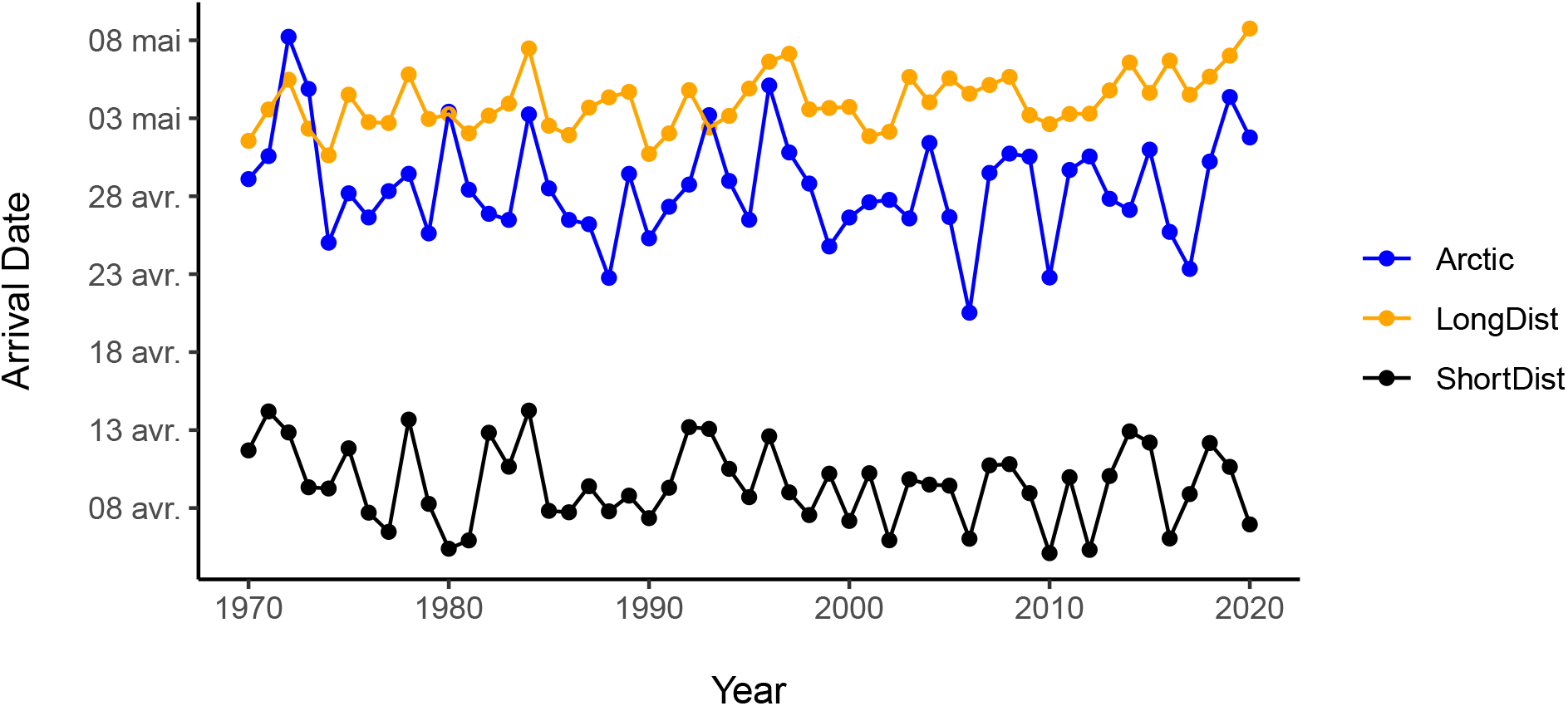
Changes in first arrival dates according to migration distance. Dates were obtained by averaging species-specific first arrival dates over 100 resamples with equal frequency.

There was a significant variation of first arrival trends among migration distances (*F* = 5.76, df = 2, 131, *p* = 0.005), with long-distance, first arrivals of non-arctic species advancing slower that those of arctic nesters and short-distance migrants (estimated marginal mean contrasts, Tukey method: *p* < 0.03 and *p* < 0.01 respectively). Trends in first arrival dates of short-distance migrants and arctic nesters did not differ (*p* = 0.01). As was the case with median arrival dates, most of the species that did advance their first arrival dates were associated to open water (Fig. 5).

**Figure 5:**
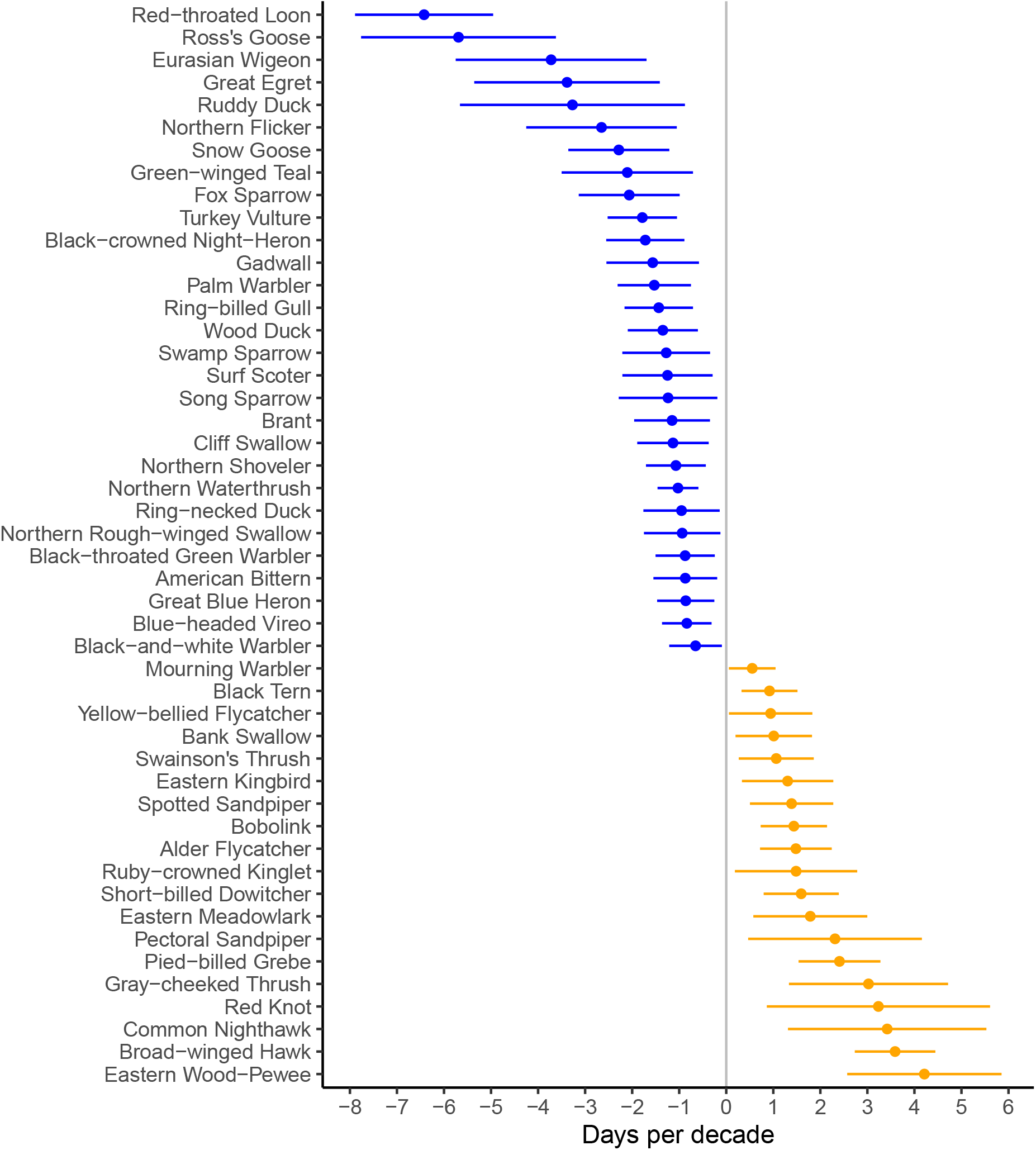
Species with significant 50-year trend in first arrival dates. Bars represent 95 % confidence intervals.

Spark’s ‘median bird’ (1980) method and our sampling procedure for first arrival dates were designed to remove correlations between trends in arrival dates and sampling effort or species abundances. As expected, we found no significant correlation between arrival dates and species trends over the last 51 years as expressed by changes in the proportion of occupied 10 km × 10 km squares (*r* < 0.1, *p* > 0.05; A. Desrochers, unpubl.data).

Species whose median arrival dates advanced between 1970 and 2020 tended to have earlier first arrival dates as well (*r* = 0.44, *p* < 0.001, *n* = 132). Despite this relationship, there was substantial variation between responses based on the two indicators (Fig. 6). First arrival dates of some uncommon waterfowl species advanced much faster than their median arrival dates but the reverse happened for some long-distance migrants such as Eastern Wood Pewee (*Contopus virens*), Common Nighthawk (*Chordeiles minor*) and Broad-winged Hawk (*Buteo platypterus*; Fig. 6).

**Figure 6:**
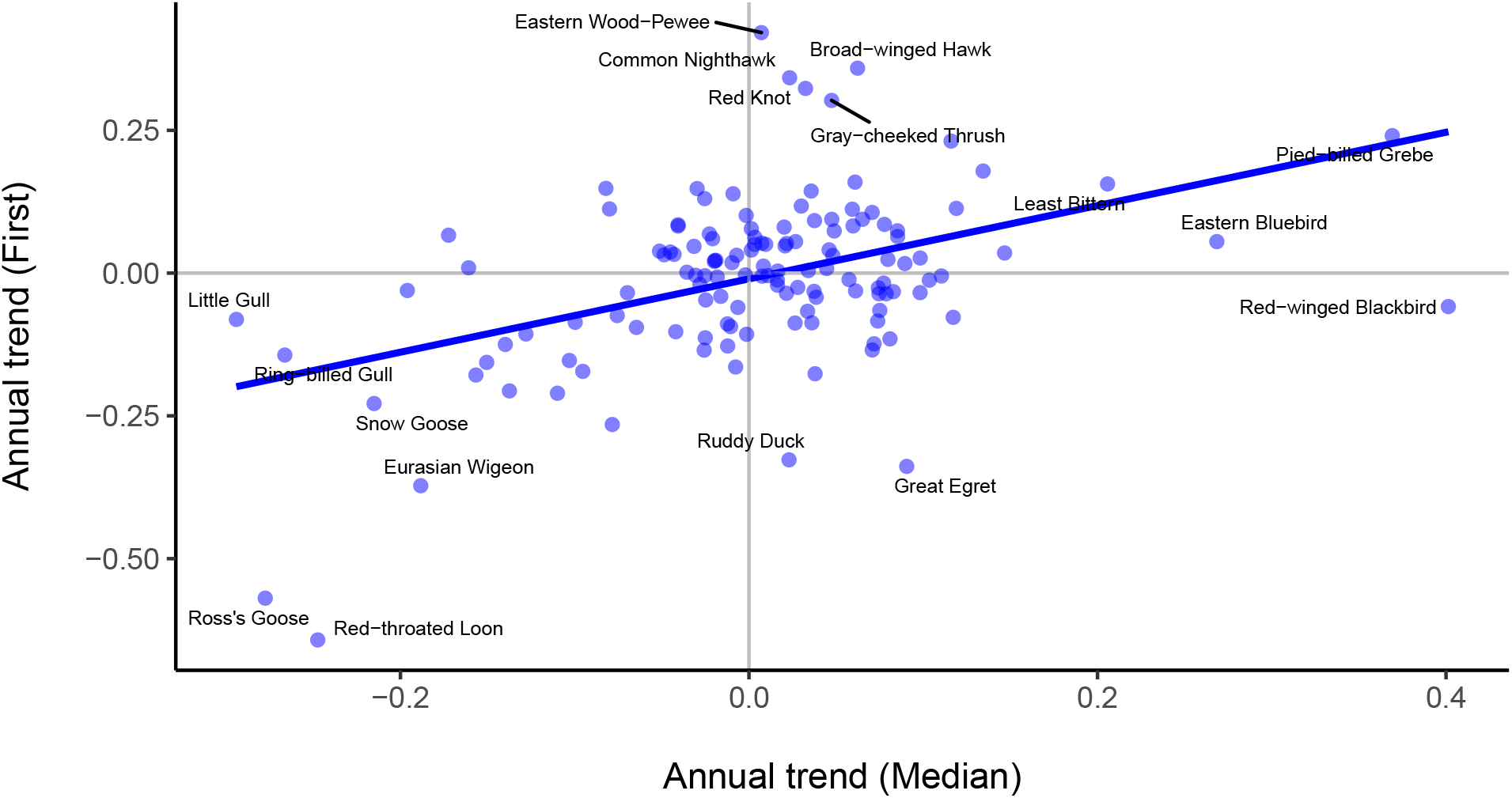
Relationship between 1970-2020 trends in first vs. median arrival dates. Each point represents one species. Outlying species identified only, for clarity.

Early arriving species showed greater interannual variation in their migration dates than later arriving species (standard-deviation vs. mean: *r* = −0.31, *p* < 0.001, *n* = 149). A similar pattern was found for first arrival dates (*r* = −0.27, *p* < 0.001, *n* = 145).

## Discussion

Shifts in spring migration dates reported in the ornithological literature are the outcome of at least three processes: actual changes in phenology, changes in abundance, and changes in sampling effort. Higher abundances or sampling effort would in most cases lead to earlier dates. Unlike most studies published on the subject, the current study was able to reduce or eliminate potential bias in arrivals due to long-term changes in abundance or sampling effort. In contrast to Horton et al.’s (2020) recent and extensive examination of North American species, we found no consistent change in the spring arrival phenology of birds in northeastern North America. However, we were able to show a great flexibility in migration dates over the last 51 years. A substantial proportion of the species examined shifted their spring arrival dates, one way or another. Among those, several ducks, larids, and arctic-nesting species have advanced their migration by as much as three weeks. By contrast, species that now tend to arrive later included many insectivore neotropical migrants, a group often considered vulnerable to the mismatch between the times of peak food supply and demand during the nestling period (Saino et al., 2011; Both et al., 2009).

### The mismeasure of migration

At first glance, documenting the phenology of spring migration would strike as a simple task, but the literature on the subject shows the opposite. Despite the large variety of data, metrics and modeling methods available, early studies of spring migration phenology often relied on arrival dates. Despite their flaws, first arrival dates have often been used and defended because they were the only available data (e.g. Rubolini et al., 2007) and possibly because of the lack of advanced statistical methods. Knudsen et al. (2007) and Lehikoinen and Sparks (2010) reviewed sampling and analytical problems encountered with the combination of noisy data such as arrival dates, and simplistic analytical approaches: truncation, missing data, autocorrelation, nonstationarity, etc. Warnings such as those by Knudsen, Lehikoinen and others are essential to the advancement of the field, but researchers should also consider the tradeoff between transparency and sophistication difficult to comprehend even for statistically-literate readers (e.g. Horton et al., 2020).

Even when sampling bias is dealt with by simple and replicable methods such as those of the present study, limitations remain. For example, our analysis, as many others, estimates the phenology en route or at stopover sites, while most birds observed were presumably en route to their breeding grounds, providing no information on the ultimate arrival dates. Furthermore, in the absence of marked individuals, most studies of temporal changes confound lifetime changes within individuals (especially among long-lived species), and selection among individuals with unchanging migration phenologies. Also, in the case of species breeding in the latitudinal range of the present study, median dates are simply those for which half of the total counts before 10 June were obtained. Thus, the actual meaning of ‘median’ arrival dates is not independent of the breeding range.

The choice of a time scale adds to the challenge of measuring migration phenology. Lehikoinen warned that “The methodological message provided by long time series is that we should be cautious when short recent time series are used, because of varying directions of ‘trends’ ” (Lehikoinen et al., 2004). Despite shorter-term fluctuations, there is evidence that six common species in Finland have gradually advanced their spring migration since the end of the Little Ice Age (1750— 1988; Fig. 2 in Lehikoinen et al., 2004). Ahas (1999) used “arrival dates” from a 132 year time series from two species (*Alauda arvensis*, *Motacilla alba*), and found that trends in arrival dates depended on the timescale used, with an overall trend toward later arrivals by those two species. Ahas (1999) did not define “arrival dates”, thus making it impossible to determine possible biases. Mason (1995) reported first arrival dates of 23 species from 1942 to 1991 in England, but did not account for possible effects of a fivefold increase in observer effort over the study period, or a possible abundance bias. Nevertheless, Mason found no overall advance in arrival dates in the 1942—1991 period and speculates that observers ‘have always been assiduous in noting [bird arrivals]’. Unfortunately, an absence of change in ‘assiduity’ does not remove the bias due to changing numbers of observers or birds observed.

### A Nearctic perspective

How similar are trends in spring migration in eastern North America relative to Europe and Asia? Based on available literature at the time, Knudsen et al. (2011) concluded that changes in migration phenology were similar between Palearctic and Nearctic birds. However, North American studies published so far were of relatively short duration, possibly leading to unreliable estimates. Hurlbert et al. (2012) argued that in the United States, southeastern species advanced their migration dates more than northern counterparts, over the period 2000-2010. But more recently, Horton et al. (2020) concluded from weather radar data that birds in the United States substantially advanced migration dates from 1995 to 2018, especially in the north of the country. Horton et al.’s findings are consistent with other studies suggesting that advancement of spring migration has been stronger in boreal latitudes than in more southerly latitudes (Rubolini et al., 2007; Horton et al., 2020; Post et al., 2018; but see Parmesan, 2007). If estimates from Horton et al. (2020) as well as Palearctic studies (Lehikoinen et al., 2004; Rubolini et al., 2007) are to be trusted, we would expect a general advance of spring migration by one to three weeks over the last 51 years. Those estimates are inconsistent with what Quebec birders have experienced and reported in the same period, as shown in the present study.

An earlier, unpublished, study based on ÉPOQ concluded that the spring migration of 113 species advanced in Quebec over the period 1969-2008, by 1.3 day per decade (Francoeur, 2012), a figure consistent with the 1.0 day per decade obtained by Lehikoinen et al. (2004). However, this conclusion was based on one of nine metrics used, the 25th percentile arrival date. We question this choice of metric, given that two of the metrics available were unbiased (first arrival and median dates based on random, equal-sized samples) and yielded much subtler trends in migration dates. Constrasting results among studies on north American birds migration phenology raises questions about this general advancement pattern being a ‘flagship example of the biological impacts of climate change’ as claimed by many, like Kelly et al. (2017).

### Searching for mechanisms

Widely cited studies describing phenological changes in taxa (e.g., Root et al., 2005; Parmesan, 2007) usually come with numerous caveats, but they are often cited to support claims that changes in phenology are common, unidirectional, and essentially caused by a changing climate (e.g., Kelly et al., 2017; Miller-Rushing et al., 2008; Newson et al., 2016; Settele et al., 2014). But several authors argue that we need a more ‘mechanistic’ approach to the study of migration phenology to enable attribution to climate change (Knudsen et al., 2011; Chmura et al., 2019; Thackeray et al., 2016). Without a mechanistic understanding, projections into the future will be unreliable, if not misleading. To materialize their pledge for a more ‘mechanistic’ approach, Thackeray et al. (2016) analysed 10,003 terrestrial and aquatic phenological data sets from 812 taxa, and concluded that the phenology of species at the “primary consumer” trophic level is more sensitive to climate change than that of other species.

Our theoretical understanding, and thus our expectations, about the phenotypical responses of birds to a warming climate thus remains superficial, and would benefit from more specific predictions, such as those that could emerge from regional comparisons in phenological responses. For example, if the rate of climate warming has an effect on migration phenology, we should expect a weaker phenological trend in eastern North America than what has been found in the Palearctic, because warming has been slower in eastern North America than in other regions where most of the studies on migration phenology have occurred (Fig.1 in Hansen et al. (2006)). Furthermore, arctic-nesting species should advance their migration dates more than other species, because in recent decades, global warming has been mostly concentrated in higher latitudes (IPCC, 2013). Our results concerning migration distances are consistent with the latter two predictions, as well as the findings of several other studies (Lehikoinen and Sparks, 2010), but exceptions remain (Jonzén et al., 2006; Sullivan et al., 2016).

One way by which the present study may advance our understanding of mechanisms leading to phenological changes is finding that species associated to open water, which should respond strongly to ice cover advanced their migration dates more than most other species. Butler (2003) also observed that aquatic species were among the most responsive species. Coincidentally since 1970, ice cover in early spring has retreated, from the Great Lakes to the Gulf of the St. Lawrence (Fig. 7). Further study of year-to-year variations in ice cover and other regional phenomena such as the North Atlantic Oscillation (Stenseth et al., 2003; Haest et al., 2018) and their correlation with spring arrival dates would help evaluate causal factors. A spatially-explicit examination of eBird data such as those used in the present study, and short-term weather fluctuations may provide answers, but this is outside the scope of the current study.

**Figure 7:**
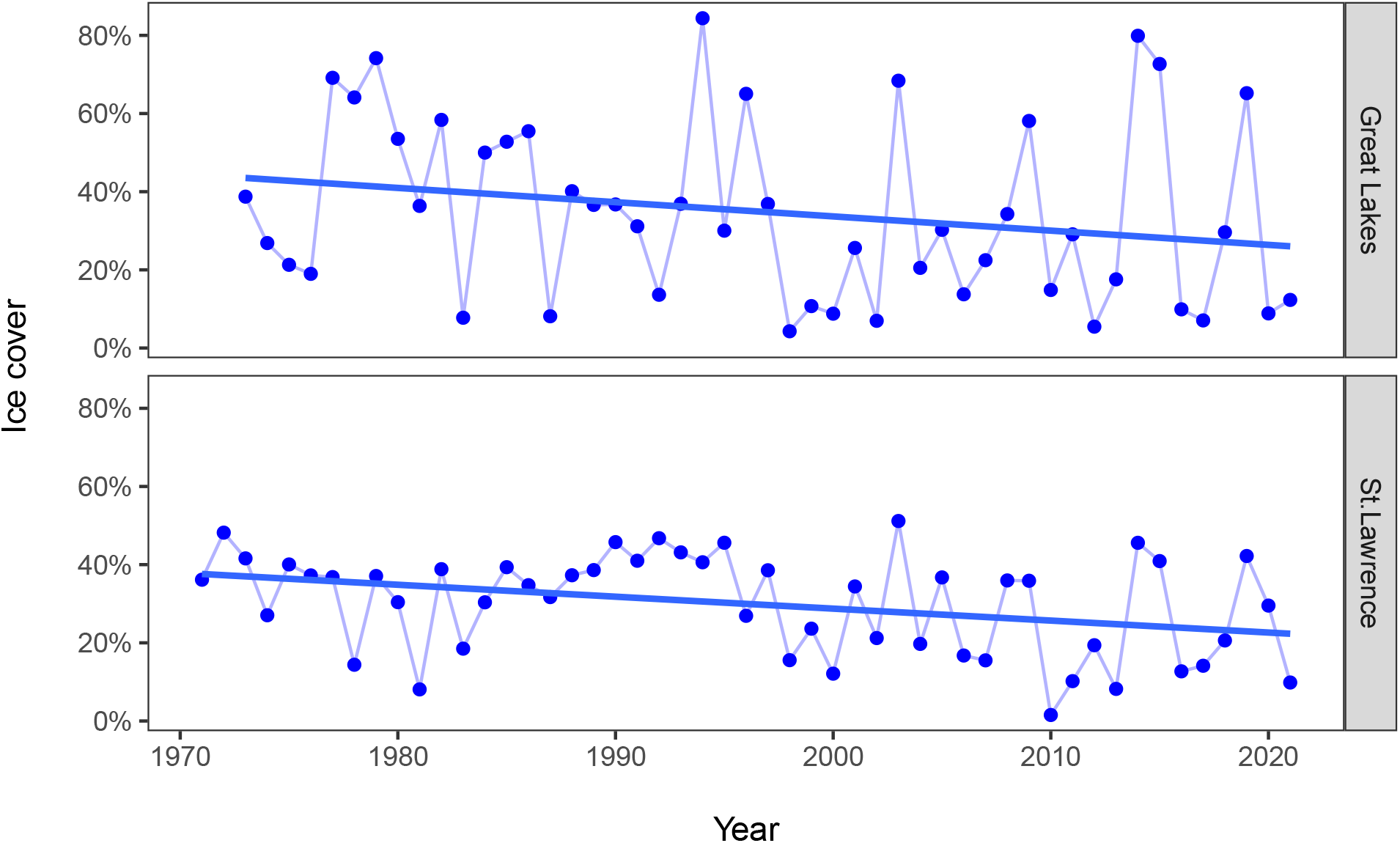
Decreasing total ice cover of the St. Lawrence river system, from the Great Lakes to the Gulf, on the first week of March. Trends were significant (*p* = 0.009) and similar between regions (*p* = 0.16). Data from Environment anc Climate Change Canada (https://iceweb1.cis.ec.gc.ca/IceGraph/)

We expect short-distance migrants to be influenced mostly by regional cues, in contrast to long-distance migrants that are driven mostly by photoperiodic and endogenous cues (Chmura et al., 2019). In several papers (e.g., Both and Visser, 2001), Both and coworkers portrayed arrival dates as relatively inflexible, especially in long-distance migrants, due to the lack of relevant cues in the wintering grounds and the apparent reliance on photoperiod (Gwinner and Helm, 2003).

Some authors argue that early migrating species are more variable in their arrival dates than later-arriving species, as was the case here, because of greater variability in early spring weather (Sparks et al., 2001). If regional weather is used as a cue by short-distance migrants, authors have hypothesized that changes in migration phenology should be greatest in short-distance migrants. However, in their review, Knudsen et al. (2011) concluded that evidence for differential trends in spring migration phenology according to migration distance was mixed. The evidence from the present study adds another layer of mixed evidence: long-distance migrants did not advance their first arrival dates as much as short-distance migrants or arctic-nesting species, but we found no significant difference among species groups when median arrival dates were used.

### Concerns about phenology

With the accumulating evidence on changes in the phenology of plants and insects in temperate and northern regions, concerns have been raised about a possible mismatch between dates of maximal food availability and dates of maximal nestling growth (Saino et al., 2011; Both et al., 2009), and its consequences on populations. Fueling this concern, Møller concluded that populations of migratory bird species that did not show a phenological response to climate change are declining (Møller et al., 2008). Visser and Both (2005) went further and argued that desynchronization may lead to ‘severe ecological dysfunction’. Those concerns are based on the possibility that arrival dates are generally not flexible enough (Both and Visser, 2001), which appears counter to the available evidence (Knudsen et al., 2011), including our own study. It remains possible that changes in migration dates are outpaced by phenological changes in breeding ranges, as has been documented in Europe (Saino et al., 2011), but the population consequences of ecological mismatches between arrival dates and optimal conditions for nesting remain little understood (Knudsen et al., 2011).

If ecological mismatch were to lead to ‘ecological dysfunction’ (Visser and Both, 2005), one would expect measurable evolutionary responses, given that factors triggering the onset of migration have a strong genetic basis (Berthold et al., 2003). Phenotypic plasticity, the facultative change in behavior or physiology induced by environmental change, is often assumed to trump Darwinian evolution as a mechanism of change in migration phenology (Van Buskirk et al., 2012; Gordo, 2007). Phenotypic plasticity of the timing of migration has been demonstrated (Saino et al., 2004; Gienapp et al., 2007), but longitudinal studies using artificial selection (Pulido et al., 2001) and passive studies based on marked individuals (Gill et al., 2014; Adriaensen et al., 1993; Møller, 2004; Brown and Brown, 2000) suggest that rapid evolution also takes place. To date, the relative roles of phenotypic vs. evolutionary responses remain poorly understood (Knudsen et al., 2011; Pulido and Berthold, 2004; Berteaux et al., 2004), with indirect evidence for changes in allele frequencies responses to changing spring climate (Jonzén et al., 2006). However, other studies came to different conclusions (Both, 2007; Jonzén et al., 2007; Gill et al., 2014). Some researchers believe that rapid evolution may not be sufficient to compensate for shifting climate patterns (Lehikoinen and Sparks, 2010; Radchuk et al., 2019). Others (Both and te Marvelde, 2007) acknowledge that evolutionary pressures for an optimal match between arrival dates and the production of food will modify departure rules based on photoperiod and may therefore answer concerns about negative impacts of changes in ecosystem vs. migration phenology.

### Conclusion

Several authors claim that the advance in spring migration is among the best documented biological responses to climate change (e.g. Miller-Rushing et al., 2008), but our study and several others question this perception. Phenological responses by migrating birds are too diverse to be the result of a simple, overarching, phenomenon. Furthermore, even if a substantial number of species have shown their ability to shift their migration phenology, many of them have not. Those birds may not experience enough pressure to change, or they may be incapable to respond swiftly to changes in climate or other phenomena occurring in spring (Both and Visser, 2001). Determining whether temporal changes in phenology, or the lack thereof, are adaptive will continue to challenge our understanding of causes and consequences of migration phenology, especially while our understanding of the patterns themselves remains unsettled.

## Acknowledgements

Thanks to Marshall Illiff and Jenna Curtis at eBird for the support with data management. Thanks to Dominique Berteaux, Jacques Larivée, Jean-Pierre Savard, and Pascal Côté for thoughtful comments on an earlier draft of this paper.

## Appendix

**Table 1:**
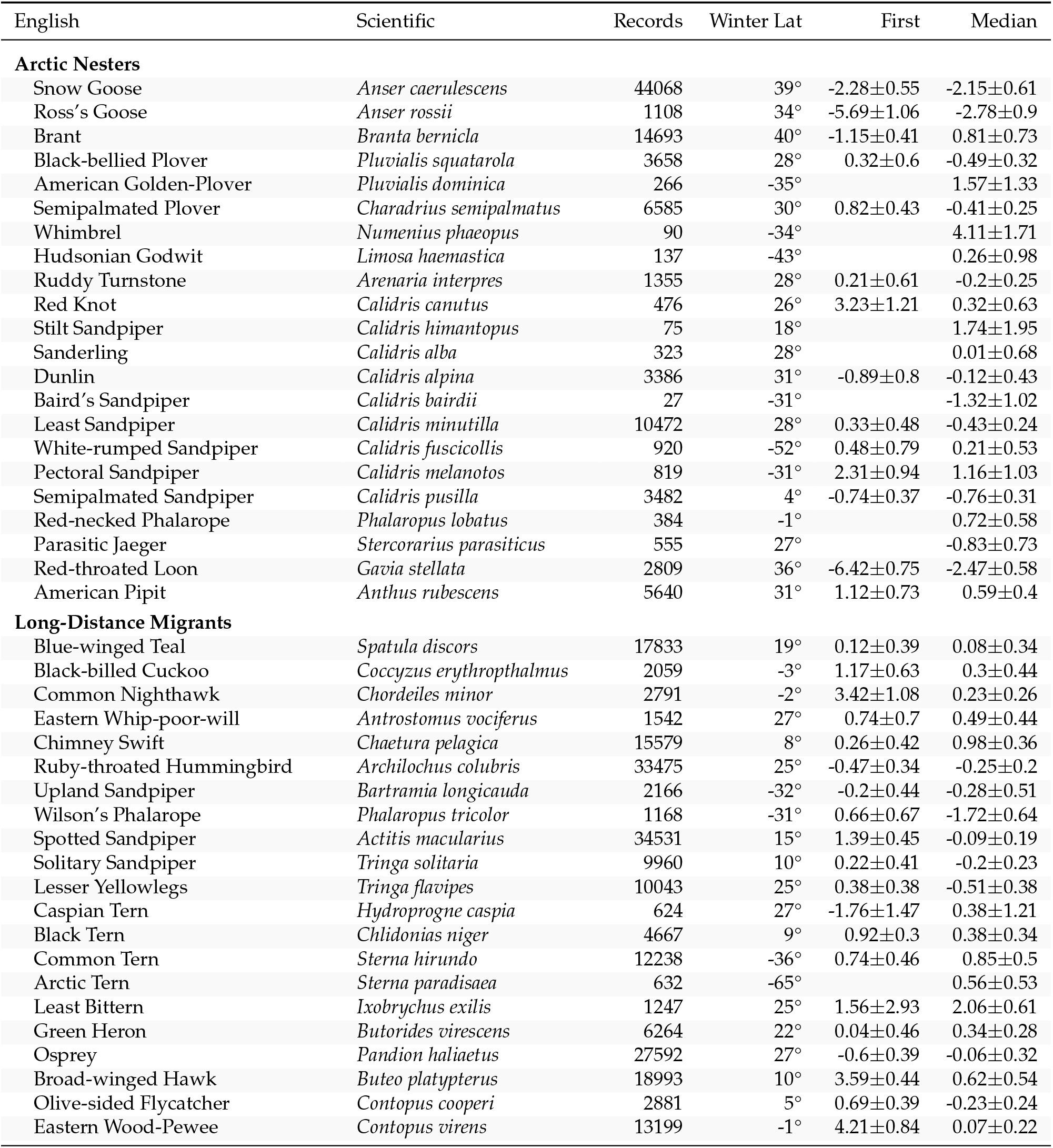
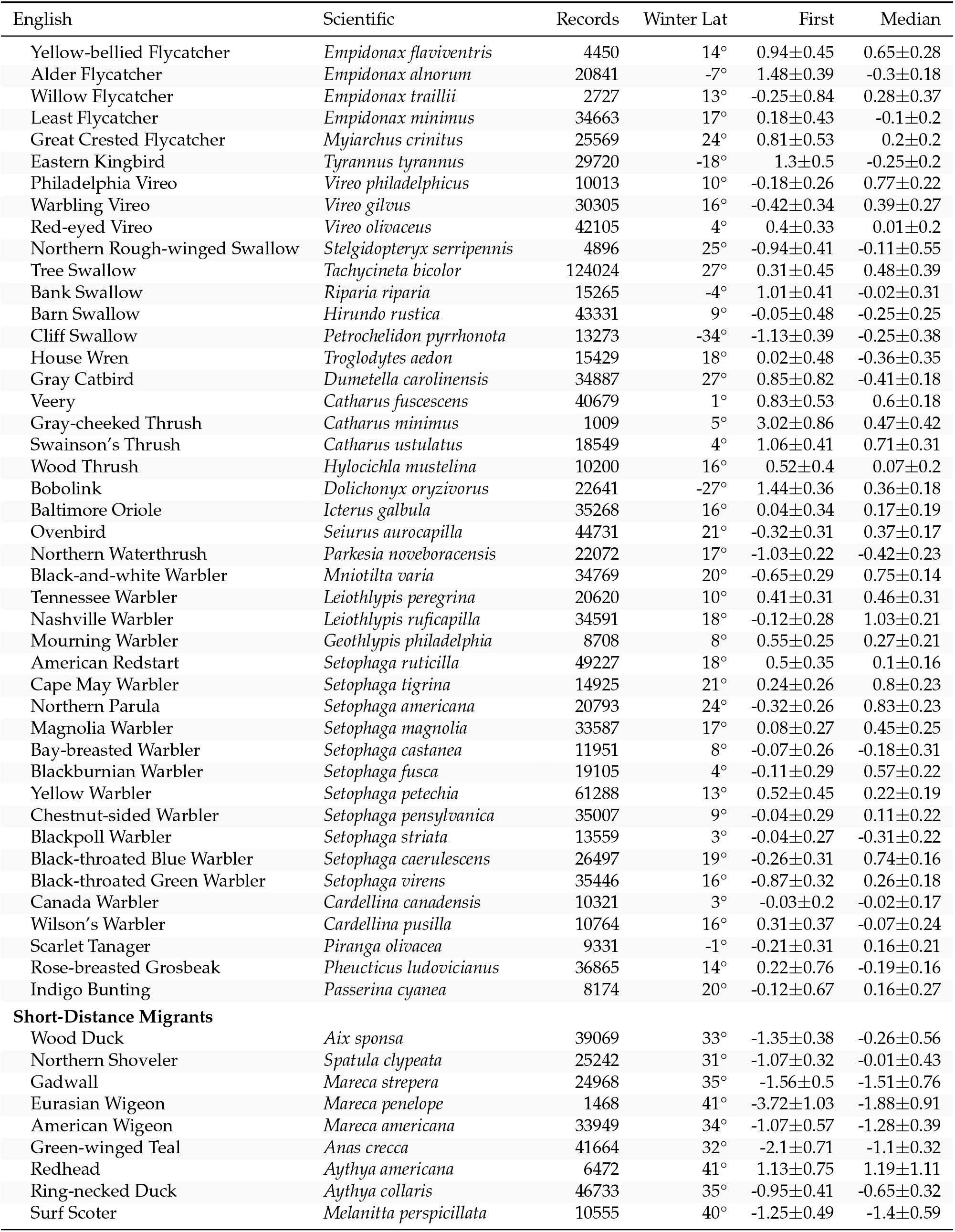
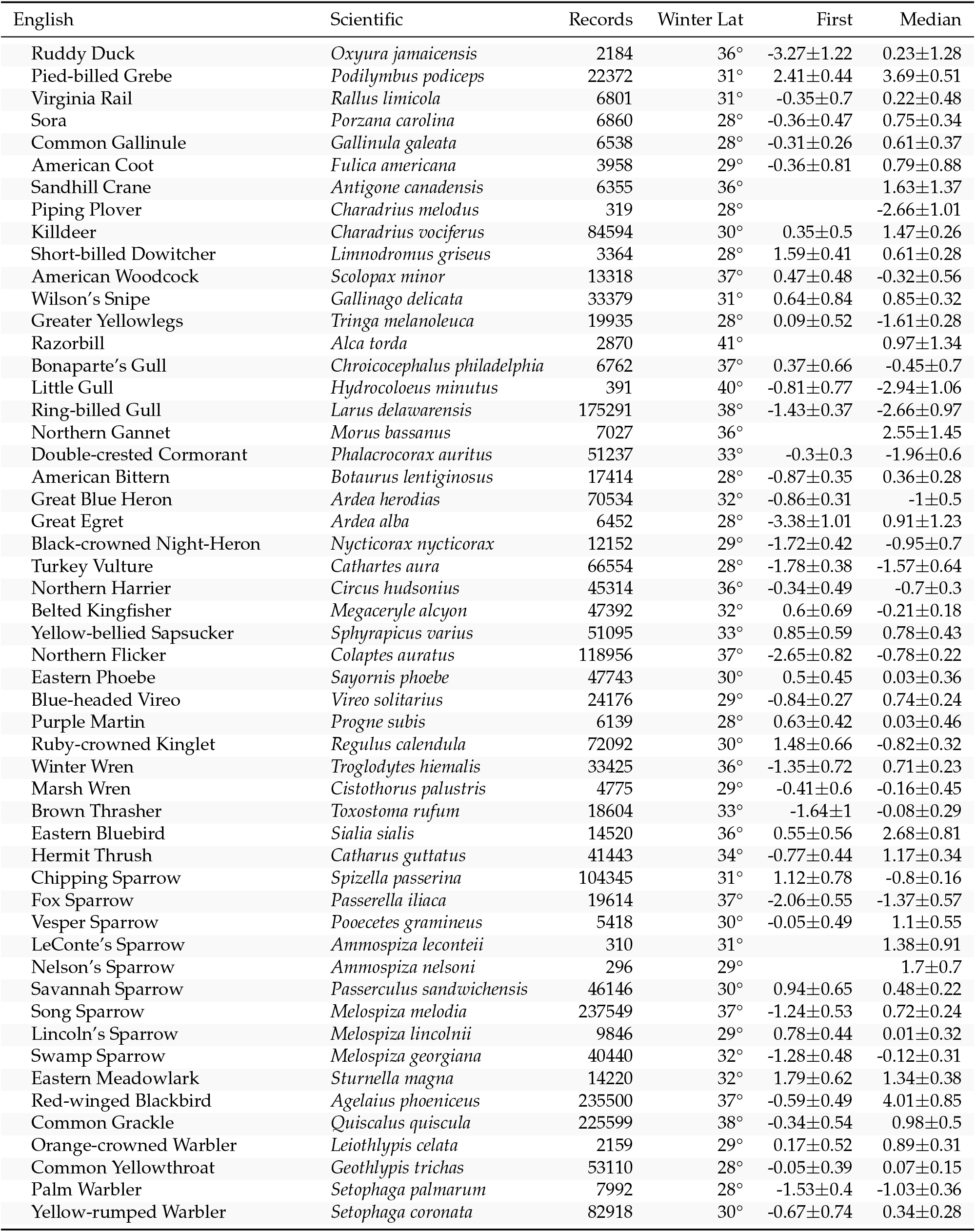
List of species, sorted by taxonomic order, within migration distance group. Records denote the number of checklist reporting the species. Latitudes are means, weighted by number of records. Means and standard errors of trend estimates (days per decade) are shown for first and median arrival dates. First arrival trends were not calculated for species with lowest annual numbers of records < 10.

## Notes

### Competing Interest Statement

The authors have declared no competing interest.

